# Describing Inhibitor Specificity for the Amino Acid Transporter LAT1 from Metainference Simulations

**DOI:** 10.1101/2022.05.03.490502

**Authors:** Keino Hutchinson, Dina Buitrago Silva, Joshua Bohlke, Chase Clausen, Allen A. Thomas, Massimiliano Bonomi, Avner Schlessinger

**Affiliations:** Department of Pharmacological Sciences, Icahn School of Medicine at Mount Sinai, New York, NY 10029; Department of Bioengineering and Therapeutic Sciences University of California, San Francisco, San Francisco, CA 94143; Department of Chemistry, University of Nebraska at Kearney, Kearney, NE 68849; Institut Pasteur, Université Paris Cité, CNRS UMR 3528, Department of Structural Biology and Chemistry, 75015 Paris, France

## Abstract

The human L-type amino acid transporter 1 (LAT1; SLC7A5) is a membrane transporter of amino acids, thyroid hormones, and drugs such as the Parkinson’s disease drug L-Dopa. LAT1 is found in the blood-brain-barrier (BBB), testis, bone marrow, and placenta, and its dysregulation has been associated with various neurological diseases such as autism and epilepsy as well as cancer. In this study, we combine metainference molecular dynamics (MD) simulations, molecular docking, and experimental testing, to characterize LAT1-inhibitor interactions. We first conducted a series of molecular docking experiments to identify the most relevant interactions between LAT1’s substrate binding site and ligan ds, including both inhibitors and substrates. We then performed metainference MD simulations using cryo-EM structures in different conformations of LAT1 with the electron density map as a spatial restraint, to explore the inherent heterogeneity in the structures. We analyzed the LAT1 substrate binding site to map important LAT1-ligand interactions as well as newly described druggable pockets. Finally, this analysis guided the discovery of previously unknown LAT1 ligands using virtual screening and cellular uptake experiments. Our results improve our understanding of LAT1-inhibitor recognition, providing a framework for rational design of future lead compounds targeting this key drug target.

**Statement of Significance:** LAT1 is a membrane transporter of amino acids, thyroid hormones, and therapeutic drugs, that is primarily found in the BBB and placenta, as well as in tumor cells of several cancer types. We combine metainference MD simulations, molecular docking, and experimental testing, to characterize LAT1-inhibitor interactions. Our computational analysis predicts S66, G67, F252, G255, Y259, W405 are critical residues for inhibitor binding and druggable sub-pockets in the outward-occluded conformation that are ideal for LAT1 inhibitor discovery. Using virtual screening and functional testing, we discovered multiple LAT1 inhibitors with diverse scaffolds and binding modes. Our results improve our understanding of LAT1’s structure and function, providing a framework for development of future therapeutics targeting LAT1 and other SLC transporters.

## INTRODUCTION

The L-type amino acid transporter 1 (LAT1; SLC7A5) mediates the import of large neutral amino acids such as phenylalanine and tyrosine, in a 1:1 exchange for intracellular amino acids (e.g., glutamine)^1^. LAT1 is a member of the Amino acid, polyamine, and organocation (APC) superfamily that includes Na^+^ independent secondary transporters and plays an important role in a broad range of biological processes^73^. LAT1 is among the most abundant transporters in the blood-brain-barrier (BBB), with 100-fold greater expression levels in BBB endothelial cells relative to other tissues^2^. LAT1 can also be found in the placenta^3^, where it delivers essential neutral amino acids and thyroid hormones, making it a key protein involved in cell growth and development. For example, leucine is imported by LAT1 to activate the mechanistic target of rapamycin complex 1 (mTORC1), a known regulator of cell growth and metabolism^4,5^.

Genetic variations of LAT1 have been associated with multiple diseases and disorders. LAT1 has recently been associated with epileptic seizures due to its role in negatively regulating the ion channel Kv1.245^18^. Additionally, rare mutations in coding regions of the *SLC7A5* gene (A246V and P375L), have been associated with autism spectrum disorder (ASD)^19^. These mutations decrease the activation of the amino acid response (AAR) pathway with a corresponding reduction in mRNA translation, thereby leading to an adverse outcome of neuronal activity, microcephaly and other neurobehavioral problems related to ASD^19^. LAT1 overexpression has been linked to disease in numerous cancer types^20–29^, particularly in prostate^21^, gastric^22^, and pancreatic^23^ cancers. This upregulation of LAT1 increases the activation of the mTOR pathway, leading to the hyperproliferation of cells in the immune system and cancer^30^.

In addition, LAT1 is highly important pharmacologically. It transports drug^31–35^ and prodrug^36–41^ substrates that are delivered specifically into cells or tissues expressing this transporter. For example, LAT1 mediates the transport of the CNS drugs gabapentin^31^, L-DOPA^38^, pregabalin^32^, and melphalan^33^ across the BBB. LAT1 is also an emerging therapeutic target for cancer, where LAT1 inhibition deprives tumor cells from nutrients that fuel cancer cell proliferation^20^. For instance, the LAT1 inhibitor JPH-203 is currently under clinical investigation for treating Cholangiocarcinoma^34^. Furthermore, due to its increased expression of LAT1 in T-cells, it can also be targeted to augment cancer immunotherapy treatment as well as for treating auto immune diseases^29^. Other LAT1 inhibitors that have been reported are KMH-233^41^ (a phenylalanine derivative with IC_50_=18 μM), 3-iodo-L-tyrosine^46^ (a tyrosine derivative with IC_50_= 7.9 μM) and *meta*-substituted phenylalanine derivatives^43,74^ (IC_50_ = 5-10 μM) than contain large lipophilic moieties.

Despite its biological importance, our understanding of LAT1 substrate and inhibitor specificity is still lacking^11^. We have previously developed structure-activity relationship (SAR) models that allowed us to predict whether compounds are more likely to be a substrate or an inhibitor, which is highly relevant to drug delivery applications^42–46^. For example, we discovered that LAT1 is considerably more tolerant of structural changes to its ligands than was previously thought^46^. LAT1 does not require a carboxylic acid to be present in the substrate^42^, and substitution with various polar functional groups that could be used in linking amino acids to drug molecules is viable^43^. In contrast, substitution with larger groups led to potent inhibitors^74^.

LAT1 forms a heterodimer complex with type-II membrane glycoprotein 4F2 cell-surface antigen heavy chain (4F2hc; SLC3A2)^6^. While SLC3A2 does not play a specific role in transport, it aids in localization to the plasma membrane and improves the stability of transport by LAT1^7^. Recently, two independent groups determined complex LAT1/4F2hc structures in both outward-occluded and inward-open conformations^8,9,10^. These structures revealed that LAT1 has 12 transmembrane helices (TMs), with 10 core TMs that follow a LeuT-like fold as seen in other APC members, where the TMs are arranged in a 5+5 inverted repeats topology. The structures also confirmed the putative substrate-binding site and modes of substrate recognition proposed by homology models based on prokaryotic homologs^11,12^. Similar to related transporters^13^, LAT1 uses a gated-pore mechanism, adopting different conformations during transport. Structures of homologs of LAT1, including the arginine/agmatine antiporter (AdiC)^14,15^ from *Escherichia coli* and Amino acid, polyamine and organocation transporter^16,17^ (GkApcT) from *Geobacillus kaustophilus* have revealed potential conformations, providing templates to characterize LAT1 in various stages of the transport cycle.

In this study, we used various computational and experimental approaches to improve our understanding of the structural basis for LAT1 interaction with inhibitors. We first performed molecular docking to determine the most relevant interactions between the LAT1 and its inhibitors. Next, we conducted metainference MD simulations, which allowed for evaluating the inherent heterogeneity in the recently published LAT1 cryo-EM structures and potential ligand binding modes. The proposed specificity determinants were then tested by the prediction of putative LAT1 inhibitors and experimental testing of top scoring hits using a cell-based assay. Finally, we discuss these results and how they can be used to describe mechanisms of ligand binding and transport by LAT1, as well as the relevance of our approach to study other Solute Carrier (SLC) transporters.

## MATERIALS AND METHODS

### Molecular docking

Molecular Docking was performed using Glide^47^ from the Schrödinger suite. All ligands were docked to the recently solved cryo-EM structures of LAT1 (PDB: 6IRT^8^, 7DSL^10^). These structures were prepared for docking with the Glide Protein Preparation under default parameters. The ligand binding site was defined based on the coordinates of the respective ligand in each published structure. The receptor grid for docking was generated *via* Glide Receptor Grid Generation panel. The small molecules used in molecular docking were prepared for docking using LigPrep of the Schrödinger suite using default parameters. The docking results were visualized via PyMOL^48^.

### Enrichment analysis

Docking and enrichment calculations were performed with Glide as described above. 8 known ligands of LAT1 were selected from the literature, using available IC_50_ data and ability to dock into the inward-open conformation as selection criteria. 635 decoys were then generated with the DUD-E server^49^ using these ligands. This set of 643 molecules was then screened against both inward-open (6IRT) and outward-occluded (7DSL) structurers, at the substrate binding site for LAT1. The results were evaluated by calculating the AUC (Area Under the Curve) and LogAUC, representing the ability of the virtual screening to discriminate known ligands among the set of decoys^64,65^. The corresponding plots (Fig. 1) show the percentage of known ligands correctly predicted (y-axis) within the top ranked subset of all database compounds (x-axis on logarithmic scale).

**Figure 1:**
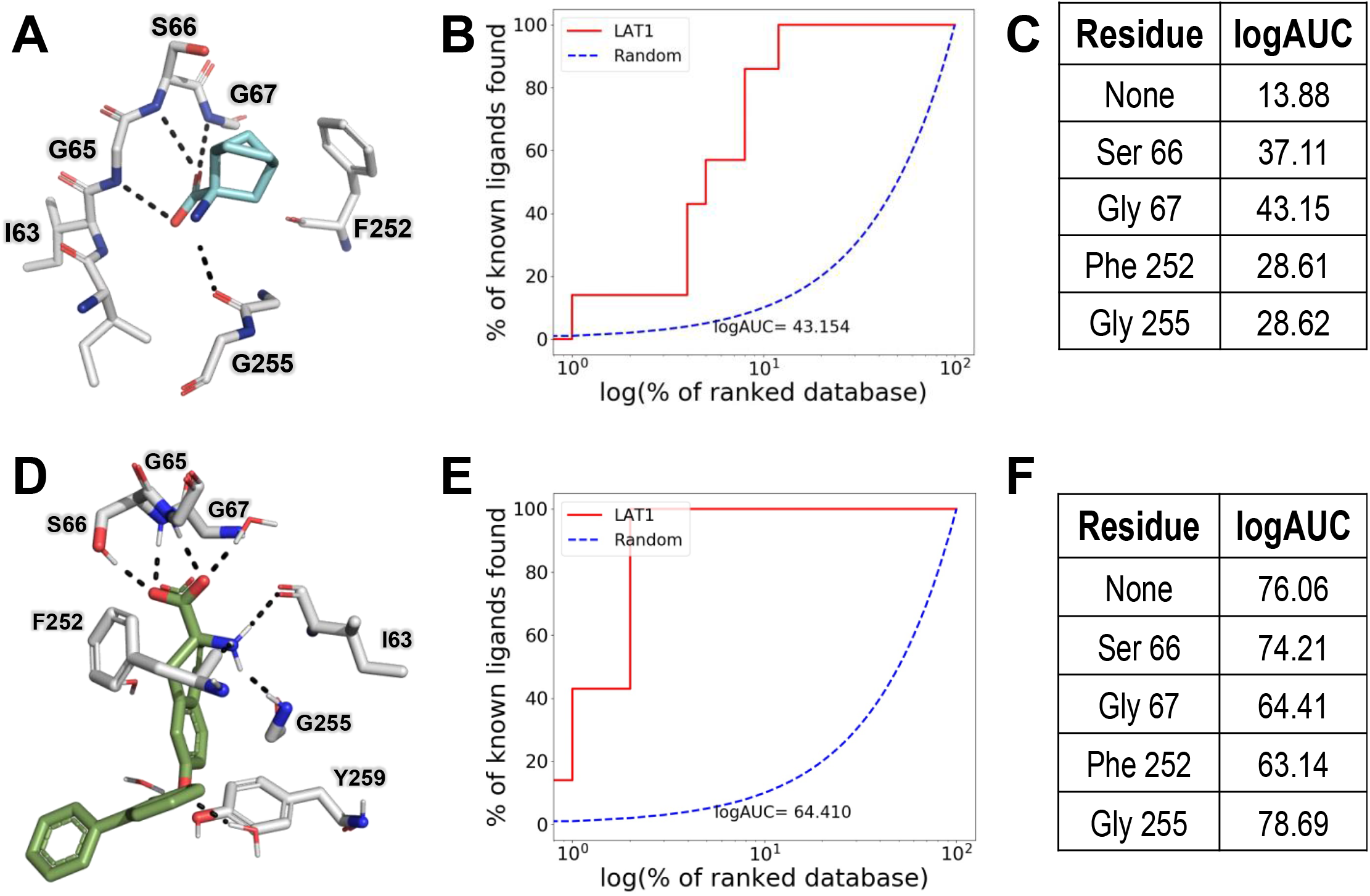
Ligand enrichment for LAT1 structures. Key binding site residues are shown as sticks where oxygen, nitrogen, and carbon atoms are represented in red, blue, and white (A) or white (B-C), respectively. Hydrogen bonds are displayed as a dashed black line. (**A**) Docking pose of BCH to LAT1 guided by enrichment. BCH is predicted to form hydrogen bonds with G65, S66, and G67, key residues for LAT1 amino acid recognition. (**B)** The logarithmic area under the curve (logAUC) derived from constrained docking at residue G67 (red line; logAUC=64.4), as compared to random selection of ligands from a set of ligands and decoys (dashed blue line); **(C)** Table showing logAUC values for constrained docking to selected residues of LAT1 substrate binding site in the inward-open conformation (**D**) Binding pose of JX-078 to LAT1 found in published outward-occluded structure **(E)** logAUC values of constrained docking to residue G67 **(F)** logAUC values for constrained docking to selected residues of LAT1 substrate binding site in the outward-occluded conformation.

### Virtual Ligand Screening

We used Glide to virtually screen the ZINC20^50^ Lead-Like library (‘In Stock’) and Enamine amino acid library against the two LAT1 conformations. We then selected compounds that contained an amino acid group from the 1,000 top-scoring compounds of the Lead-Like library screen and did the same for the Enamine screen, as it contained many amino acid derivatives. We prioritized compounds predicted to interact with the binding site through hydrogen bonding with important residues (Fig. 1) and removed compounds with an unlikely pose or strained conformation^66–68^.

### Pocket Volume Measurement

POVME3^51^ (Pocket Volume Measurer 3) was used to calculate binding site volumes. We used default parameters for Ligand-defined inclusion region, using BCH (PDB:6IRT) and JX-078 (PDB:7DSL) as the reference ligands, respectively.

### Metainference MD simulations

The coordinates of LAT1 (chain B) were extracted from the PDB structures 6IRT, 7DSL cryo-EM structures, including the respective ligands (BCH, JX-078). CHIMERA^52^ was used to select the cryo-EM density within 5 Å of the structure^53^, which was then used as input of the Metainference simulation of LAT1-ligand binding. The starting model was prepared using CHARMM-GUI^54^ 92 POPC lipids were added to the system along with 13,297 water molecules in a triclinic periodic box of volume equal to 726 nm^3^. 31 potassium and 30 chloride ions were added to ensure charge neutrality and a salt concentration of 0.15 M. The CHARMM36m force field^56^ was used for the protein, lipids, and ions; the CHARMM General Force Field and the TIP3P model were used for the ligand and water molecules, respectively. A 30 ns-long equilibration was performed using GROMACS^57^ following the standard CHARMM-GUI protocol consisting in multiple consecutive simulations in the NVT and NPT ensembles. During these equilibration steps, harmonic restraints on the positions of the lipids, ligand, and protein atoms were gradually switched off.

In the metainference simulation, a Gaussian noise model with one error parameter for each voxel of the cryo-EM map was used. 16 replicas of the system were used, and their initial configurations were randomly selected from the last 10ns-long step of the equilibration protocol. The metainference run was conducted for a total aggregated time of 10.8 μs for 6IRT, 30 μs for the 7DSL and 20.5 μs for 7DSK. All simulations were carried out using GROMACS 2020.1 and the ISDB module^81^ of the open-source, community-developed PLUMED library^80^ [GitHub ISDB branch; https://github.com/plumed/plumed2/tree/isdb]. For the analysis, the initial frames of the trajectory of each replica, corresponding to 20% of the total simulation time, were considered as additional equilibration steps under the cryo-EM restraint and discarded. The remaining conformations from all replicas were merged and clustered using: i) the Root Mean Square Deviation (RMSD) calculated on all the heavy atoms of the ligand and of the protein residues within 5 Å of the ligand in at least one member of the ensemble; ii) the gromos algorithm^70^ with a cutoff equal to 1.0, 1.5 and 2.0 Å.

### Metainference MD Simulation Trajectory Analysis

Root mean square deviation (RMSD) and root mean square fluctuation (RMSF) were calculated using MDAnalysis^69^. RMSF was calculated using the initial frame as the reference frame and all frames were aligned to the receptor backbone. The LAT1-JX-078 interaction map was generated using ProLif^84^.

### Data availability

The GROMACS topology and input files as well as PLUMED input files used are available on PLUMED-NEST (https://www.plumed-nest.org/) under accession ID plumID:22.018.

Scripts used for post metainference trajectory analysis are available at: https://github.com/schlessinger-lab/LAT1-metainference

### Cell Culture Preparation of hLAT1 Stable Cell Line

TREx HEK-hLAT1 (XenoPort Inc., Santa Clara, CA)^74,85^ is an inducible cell line that is under the control of a tetracycline inducible promoter for LAT1 and a secondary tetracycline inducible plasmid for 4F2hc (SLC3A2). Both LAT1 and 4F2hc are important components for uptake activity, as they form a heterodimer to promote native transport of their substrates^6^. Doxycycline (Dox) and sodium butyrate, a histone deacetylase (HDAC) inhibitor, are supplied in the cell media to promote expression of LAT1 and 4F2hc 24 hours before experimentation occurs. The creation of HEK-hLAT1 stable cells is previously described in further details^11^. Cell culture preparation of these cells requires that they are grown and maintained in Dulbecco’s Modified Eagle Medium (DMEM) media with 100 units/mL penicillin, 100 units/mL streptomycin, 2 mM L-glutamine, 1 μg/mL Fungizone, 10% fetal bovine serum, and 3 μg/mL blasticidin. Cells were grown in a humidified incubator with 5% CO_2_ at a stable 37 °C.

### *cis*-Inhibition Assay and IC_50_ Computations

TREx HEK-hLAT1 cells were plated at a seeding density of 0.2 × 10^6^ cells per well in poly-D-lysine coated 24-well plates in DMEM medium described above. Cells were grown to at least 90% confluency 48 hours post seeding and 24 hours before the assay, the cells were induced with 2 mM sodium butyrate and 1 μg/mL doxycycline for proper LAT1 expression^11^. On the day of the assay, cells are washed twice with warm sodium-free choline buffer (140 mM choline chloride, 2 mM potassium chloride, 1 mM magnesium chloride, 1 mM calcium chloride, 1 M Tris) and then incubated with the same solution for 10 minutes. For the *cis*-inhibition assay, the choline buffer was removed and replaced with radioactive uptake buffer (sodium-free choline buffer containing 6 nM of [^3^H]-gabapentin) and each compound at a final concentration of 200μM. For the IC_50_ determinations, the choline buffer was replaced with radioactive uptake buffer mixed with varying concentrations of each inhibitor ranging from 0.1 μM to 200 μM. For each assay, the uptake was performed in an incubator at 37 °C for 3 minutes and the reaction was terminated by washing the cells twice with ice-cold choline buffer. The cells were then lysed with 600 μL of lysis buffer (0.1 N NaOH, 0.1% SDS) at room temperature overnight or on a shaker for 1 hour. Only 150 μL of lysis is used to measure intracellular radioactivity which is determined by scintillation counting on a LS6500 scintillation counter (Beckman Coulter Inc., Bea, CA). IC_50_ calculations were analyzed using GraphPad Prism version 9.3 (San Diego, CA) and % [^3^H]-gabapentin uptake is normalized relative to % inhibition by L-phenylalanine at 200 μM.

### Chemistry: general information

Flash chromatography purification was performed using a Teledyne ISCO Flash Chromatography System. Preparative HPLC was performed using a Gilson PLC 2020. Column: Synergi 4 μ Fusion-RP by Phenomenex, 150 × 21.2 mm. The following preparative HPLC method employed elution at 20 mL/min flow rate: A: gradient elution over 40 min of 5-40% CH_3_CN with water, both of which contained 0.1% TFA. LC-MS analysis was performed using an Agilent G6125 single quad ESI source and a 1260 Infinity HPLC system (G7112B Binary Pump and G7114A Dual λ Absorbance Detector). Column: Synergi 4μ Fusion-RP by Phenomenex, 150 × 4.6 mm. The following LC-MS method was performed using a 1.0 mL/min flow rate: A: gradient elution over 10 min of 10-80% CH_3_CN with water, both of which contained 0.1% formic acid. ^1^H NMR and ^13^C NMR were recorded on an Avance III HD Bruker instrument operating at 400 MHz and 100 MHz, respectively.

### Chemistry: synthesis

Preparation of benzyloxy-substituted tryptophan analogs (**AT182**, **AT183**, **AT184**).

#### (±)-2-Amino-3-(4-(benzyloxy)-1H-indol-3-yl)propanoic acid hydrochloride (AT182)

The title compound was synthesized in two steps using a modification of the procedure described by Blaser^75^, as follows:

##### Step A: Preparation of (±)-acetamido-3-(4-(benzyloxy)-1H-indol-3-yl)propanoic acid

To a stirred mixture of acetic anhydride (1.3 mL, 13 mmol) in acetic acid (5 mL) was added 4-benzyloxyindole (1.0 g, 4.5 mmol) and L-serine (0.47 g, 4.5 mmol; note: the L isomer was used because it was on hand, but the D isomer or racemic serine would also give the same racemic product as demonstrated by Blaser). The mixture was heated to 80 °C and stirring continued overnight. The reaction was concentrated *in vacuo* at 80 °C and carried forward into base-mediated acetyl deprotection without purification at this step.

##### Step B: Preparation of AT182

(±)-Acetamido-3-(4-(benzyloxy)-1H-indol-3-yl)propanoic acid from Step A (0.50 g, 1.4 mmol) was heated with an aqueous 15% solution of NaOH (8.1 mL, 36 mmol) in a teflon-capped glass pressure tube at 110 °C for 2 d. After cooling to rt, carefully added conc. HCl (3 mL) and filtered the resulting tan solids, rinsing with water (2 × 3 mL). Heated solids with 1:1 CH_3_CN /water (20 mL). Filtered the warm suspension to remove undissolved impurities. Chilled the mixture in a −10 °C freezer and then saturated the aqueous phase with NaCl. Sonicated the mixture to dissolve the NaCl. Separated the phases and re-extracted the aqueous with CH_3_CN (2 × 5 mL). Concentrated the CH_3_CN phases and redissolved the residue in 1:1 CH_3_CN /water (15 mL), and then syringe filtered through a 0.45 micron filter to remove smaller, undissolved particles. Purified the crude by preparative HPLC (Method A). Converted product to the HCl salt by adding 1M aq. HCl (2 mL) to the purified product and freeze drying to remove excess water and HCl. Yield: 81 mg (18%). >95% purity by LC-MS (254 nm, Method A). ^1^H NMR (CD_3_OD) δ 7.52 (d, *J* = 7 Hz, 2H), 7.38 (m, 2H), 7.32 (m, 1H), 7.04 (m, 3H), 6.64 (d, *J* = 7 Hz, 1H), 5.24 (s, 2H), 4.36 (dd, *J* = 4, 11 Hz, 1H), 3.79 (dd, *J* = 4, 14 Hz, 1H), 3.02 (dd, *J* = 11, 14 Hz, 1H). ^13^C NMR (CD_3_OD) δ 171.8, 154.4, 140.6, 138.6, 129.6, 129.0, 124.9, 123.9, 118.0, 108.4, 106.4, 101.8, 71.2, 55.2, 29.6. *m/z* (ESI-pos) M + 1 = 311.1.

#### (±)-2-Amino-3-(6-(benzyloxy)-1H-indol-3-yl)propanoic acid hydrochloride (AT183)

Prepared by the same two step procedure as **AT182**, except replacing 4-(benzyloxy)-1H-indole with 6-(benzyloxy)-1H-indole in Step A. >95% purity by LC-MS (254 nm, Method A). ^1^H NMR (CD_3_OD) δ 7.49 (d, *J* = 9 Hz, 1H), 7.44 (m, 2H), 7.35 (m, 2H), 7.29 (m, 1H), 7.10 (s, 1H), 6.99 (d, *J* = 2 Hz, 1H), 6.82 (dd, *J* = 2, 9 Hz, 1H), 5.08 (s, 2H), 4.22 (dd, *J* = 5, 8 Hz, 1H), 3.46 (dd, *J* = 5, 15 Hz, 1H), 3.29 (m, 1H, partially overlaps residual solvent peak). ^13^C NMR (CD_3_OD) δ 171.8, 156.9, 139.1, 139.0, 129.4, 128.7, 128.5, 124.5, 122.9, 119.7, 111.3, 107.9, 97.4, 71.5, 54.6, 27.7. *m/z* (ESI-pos) M + 1 = 311.1.

#### (±)-2-Amino-3-(7-(benzyloxy)-1H-indol-3-yl)propanoic acid hydrochloride (**AT184**)

Prepared by the same two step procedure as **AT182**, except replacing 4-(benzyloxy)-1H-indole with 7-(benzyloxy)-1H-indole in Step A. >95% purity by LC-MS (254 nm, Method A). ^1^H NMR ((CD_3_)_2_SO) δ 11.2 (s, 1H), 8.41 (br s, 2H), 7.55 (d, *J* = 8 Hz, 2H), 7.41 (m, 2H), 7.34, (m, 1H), 7.18, (m, 2H), 6.90 (m, 1H), 6.74 (d, *J* = 8 Hz, 1H), 5.25 (s, 2H), 4.07 (m, 1H), 3.27 (m, 2H). ^13^C NMR ((CD_3_)_2_SO) δ 170.6, 145.10, 145.05, 137.4, 128.8, 128.4, 127.7, 127.5, 126.5, 126.3, 124.7, 124.5, 119.1, 111.4, 107.3, 103.0, 69.1, 52.5, 26.1. *m/z* (ESI-pos) M + 1 = 311.1.

Preparation of benzylcarbamoyl-substituted tryptophan analogs (**AT185**, **AT186**).

#### (±)-2-Amino-3-(5-(benzylcarbamoyl)-1H-indol-3-yl)propanoic acid hydrochloride (AT185)

The title compound was prepared in five steps (A-E) as follows.

##### Step A: Preparation of (±)-3-(2-acetamido-2-carboxyethyl)-1H-indole-5-carboxylic acid

Prepared from 1H-indole-5-carboxylic acid (3.0 g, 19 mmol) using the same procedure as in **AT182**, Step A. Crude product was carried directly to the next step without purification.

##### Step B: Preparation of (±)-3-(2-amino-2-carboxyethyl)-1H-indole-5-carboxylic acid

With stirring, heated crude (±)-3-(2-acetamido-2-carboxyethyl)-1H-indole-5-carboxylic acid from Step A with 6M aq. HCl (20 mL, 120 mmol) at 100 °C overnight. Concentrated *in vacuo* using CH_3_CN to azeotrope water (5 cycles). Dried solids under high vacuum overnight. Carried crude product into Step C without purification at this step.

##### Step C: Preparation of (±)-4’-((5-carboxy-1H-indol-3-yl)methyl)-5’-oxo-9l4-boraspiro[bicyclo[3.3.1]nonane-9,2’-[1,3,2]oxazaborolidin]-3’-ium

Using a similar procedure as described by Venteicher^74^, we protected crude (±)-3-(2-amino-2-carboxyethyl)-1H-indole-5-carboxylic acid from Step B (1.0 g, 4.0 mmol) by stirring with pyridine (1.3 mL, 16 mmol) and 9-borabicyclo(3.3.1)nonane (9-BBN, 0.5 M in THF, 16 mL, 8.1 mmol) in DMF (5 mL) overnight at rt. Reaction mixture was partitioned between EtOAc (15 mL) and 1M aq. KHSO_4_ (35 mL). Separated phases, and reextracted aqueous phase with EtOAc (35 mL). Combined organic phases were washed with water (2 × 35 mL), brine (50 mL), dried (MgSO_4_), filtered and concentrated *in vacuo*. Purified crude by trituration with hot EtOAc, followed by cooling to rt and filtration. Yield: 0.50 g (34%). ^1^H NMR ((CD_3_)_2_SO) δ 12.4 (br s, 1H), 11.3 (s, 1H), 8.27 (s, 1H), 7.74 (m, 1H), 7.43 (m, 2H), 6.52 (m, 1H), 5.68 (m, 1H), 3.84 (m, 1H), 3.29 (m, 1H), 3.17 (m, 1H), 1.30-1.81 (m, 12H), 0.45 (m, 2H).

##### Step D: Preparation of (±)-4’-((5-(benzylcarbamoyl)-1H-indol-3-yl)methyl)-5’-oxo-9l4-boraspiro[bicyclo[3.3.1]nonane-9,2’-[1,3,2]oxazaborolidin]-3’-ium

9-BBN-protected (±)-3-(2-amino-2-carboxyethyl)-1H-indole-5-carboxylic acid from Step C (0.50 g, 1.4 mmol) was stirred overnight at rt with benzylamine (0.15 mL, 1.4 mmol), (*i*-Pr)_2_NEt (0.71 mL, 4.1 mmol), and 1-[bis(dimethylamino)methylene]-1H-1,2,3-triazolo[4,5-b]pyridinium 3-oxide hexafluorophosphate (HATU, 0.77 g, 2.0 mmol) in DMF (5 mL). Partitioned reaction mixture between EtOAc (20 mL) and water (20 mL). Separated phases and re-extracted aqueous with EtOAc (20 mL). Combined organic phases were washed with water (2 × 20 mL), brine (20 mL), then dried (MgSO_4_), filtered and concentrated. Crude was purified by ISCO flash chromatography, eluting with a gradient of 30% EtOAc in hexanes to 100% EtOAc over 8 min. Yield: 0.40 g (64%). ^1^H NMR ((CD_3_)_2_SO) δ 11.2 (s, 1H), 8.91 (m, 1H), 8.23 (s, 1H), 8.02 (br s, 2H), 7.71 (d, *J* = 9 Hz, 1H), 7.42 (m, 3H), 7.33 (m, 3H), 7.23 (m, 1H), 6.46 (m, 1H), 5.61 (m, 1H), 4.52 (m, 2H), 3.93 (m, 1H), 3.36 (m, 1H), 3.10 (dd, *J* = 9, 15 Hz, 1H), 1.29-1.81 (m, 12H), 0.48 (m, 2H).

##### Step E: Preparation of AT185

Using a modification of a method originally described by Poulie^76^, the BBN protecting group was removed as follows. (±)-4’-((5-(Benzylcarbamoyl)-1H-indol-3-yl)methyl)-5’-oxo-9l4-boraspiro[bicyclo[3.3.1]nonane-9,2’-[1,3,2]oxazaborolidin]-3’-ium from Step D (0.40 g, 0.87 mmol) was stirred overnight at rt with TBAF (1M in THF, 3.5 mL, 3.5 mmol) and THF (3 mL). Water (3 mL) was added and it was stirred for an additional 5 min. Acidified with formic acid (1-2 mL) and concentrated *in vacuo* at 60 °C to remove most of the THF. Dissolved the resulting residue in warmed 20% CH_3_CN in water containing 0.1% TFA (15 mL). Syringe filtered the mixture through a 0.45 micron filter to remove undissolved particles. Purified product by two successive preparative HPLC runs (Method A). Converted product to the HCl salt by adding 1M aq. HCl (2 mL) to the purified product and freeze drying to remove excess water and HCl. Yield: 97 mg (30%). >95% purity by LC-MS (220 nm, Method A). ^1^H NMR (CD_3_OD) δ 8.26 (s, 1H), 7.69 (dd, *J* = 2, 9 Hz, 1H), 7.47 (d, *J* = 9 Hz, 1H), 7.39 (m, 2H), 7.33 (m, 3H), 7.24 (m, 1H), 4.62 (s, 2H), 4.33 (dd, *J* = 5, 8 Hz, 1H), 3.55 (dd, *J* = 5, 15 Hz, 1H), 3.41 (dd, *J* = 8, 15 Hz, 1H). ^13^C NMR (CD_3_OD) δ 171.6, 171.5, 140.4, 140.3, 129.5, 128.6, 128.1, 128.0, 127.5, 126.3, 122.1, 119.7, 113.6, 109.3, 54.4, 44.7, 27.4. *m/z* (ESI-pos) M + 1 = 338.2.

#### (±)-2-Amino-3-(6-(benzylcarbamoyl)-1H-indol-3-yl)propanoic acid hydrochloride (AT186)

The title compound was prepared according to the five-step procedure for **AT185**, except replacing 1H-indole-5-carboxylic acid with 1H-indole-6-carboxylic acid in Step A. >95% purity by LC-MS (220 nm, Method A). ^1^H NMR (CD_3_OD) δ 8.00 (s, 1H), 7.69 (d, *J* = 9 Hz, 1H), 7.60 (dd, *J* = 1, 8 Hz, 1H), 7.41 (s, 1H), 7.37 (m, 2H), 7.32 (m, 2H), 7.23 (m, 1H), 4.60 (s, 2H), 4.28 (dd, *J* = 5, 7 Hz, 1H), 3.52 (dd, *J* = 5, 15 Hz, 1H), 3.40 (dd, *J* = 7, 15 Hz, 1H). ^13^C NMR (CD_3_OD) δ 171.5, 171.3, 140.4, 137.6, 131.0, 129.5, 128.8, 128.7, 128.5, 128.1, 119.3, 119.1, 112.7, 108.4, 54.5, 44.6, 27.3. *m/z* (ESI-pos) M + 1 = 338.2.

## RESULTS

### Docking of known ligands to LAT1 structures

We applied molecular docking and ligand enrichment calculations to examine the pharmacological relevance of different conformations of LAT1’s binding site and ligands, as well as key interactions between them (Fig. 1). This analysis was also conducted to evaluate the optimal parameters for rational design of new LAT1 ligands. Specifically, we analyzed the ability of molecular docking and the cryo-EM structure to distinguish between known ligands of LAT1 and decoy compounds, which have similar physical properties to the known ligands but possess different topological properties, to reduce likelihood of binding^46^. Compounds were docked against two different conformations of LAT1: an inward-open conformation bound to the inhibitor BCH (PDB identifier: 6IRT)^8^ and an outward-occluded conformation bound to the inhibitor JX-078 (PDB identifier: 7DSL)^10^.

We docked this set of known LAT1 ligands and decoy compounds against the inward-open conformation using default parameters and obtained a logAUC value of 13.88 (Fig. 1C), which corresponds to a random selection of ligands. Conversely, similar docking parameters used for docking against the outward-occluded structure yielded logAUC value of 76.06, suggesting that this conformation can capture LAT1-ligand complementarity and may yield productive virtual screening.

We then used different parameters enforcing different LAT1-ligand interactions to specific residues that have been shown^8,10,11,79^ to be important for ligand binding and/or transport (i.e., S66, G67, F252, G255) (Fig. 1). Interestingly for the inward-open conformation, constraining ligands to interact with F252, which is thought to serve as a “gate” for LAT1 ligands^8,44^ yielded significantly improved enrichment scores, further validating the importance of this residue for LAT1 binding. For the outward-occluded structure, constrained docking to S66, G67, F252, and G255 yielded comparable enrichment scores to the scores derived from unconstrained docking. Notably, it is likely that the ability of the outward-occluded conformations to enrich for inhibitors is due to its increased size as compared to the inward-open conformation, i.e., 275 Å^3^ vs. 118 Å^3^, respectively (Fig. 2), which results from the movement of the sidechain of Y259. Taken together, these results suggest that the outward-occluded conformation better captures protein-ligand complementarity and is potentially more useful for the discovery of new ligands with structure-based virtual screening.

**Figure 2:**
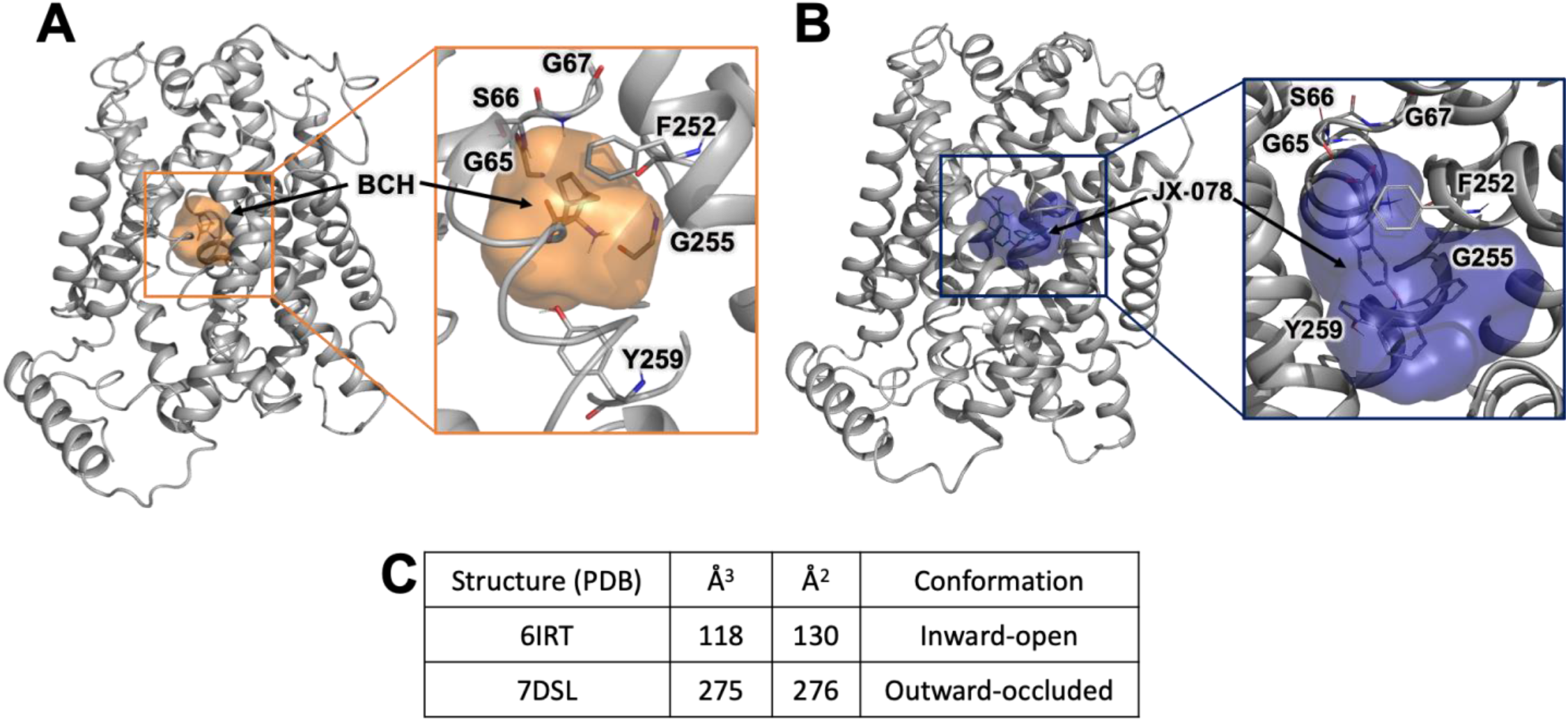
LAT1 binding site in different conformations. Visualization of LAT1 binding site and accessible regions for the **(A)** Inward-open (orange) and **(B)** Outward-occluded conformation of LAT1 (blue). **(C)** Volume and surface area of the binding site in each conformation.

### Dynamics of LAT1 binding site and inhibitor interaction

In parallel, we investigated the heterogeneity in the LAT1-inhibitor complex structure, using the metainference integrative approach^58,80^. In metainference, the molecular mechanics force field used in standard MD simulations is augmented by spatial restraints that enforce the agreement of an ensemble of replicas of the system with the cryo-EM density map^59^. Using a Bayesian inference framework, this approach accounts for the simultaneous presence of structural heterogeneity, data ensemble-averaging, and variable level of noise in different areas of the experimental map. We performed metainference simulations on the outward-occluded structure of LAT1 bound to JX-078 (7DSL). In particular, we generated clusters of conformations of the protein-ligand complex and obtained a representative model for each cluster (Fig. 3A). We assessed the stereochemical quality of each model using MolProbity^78^, which considers the number of steric clashes, rotamer outliers, and percentage of backbone Ramachandran conformations outside favored regions. Notably, all models scored better (lower) than the cryo-EM structure, with Model 1 scoring the best at 1.55 compared to the cryo-EM structure at 1.92 (Fig. S1).

**Figure 3:**
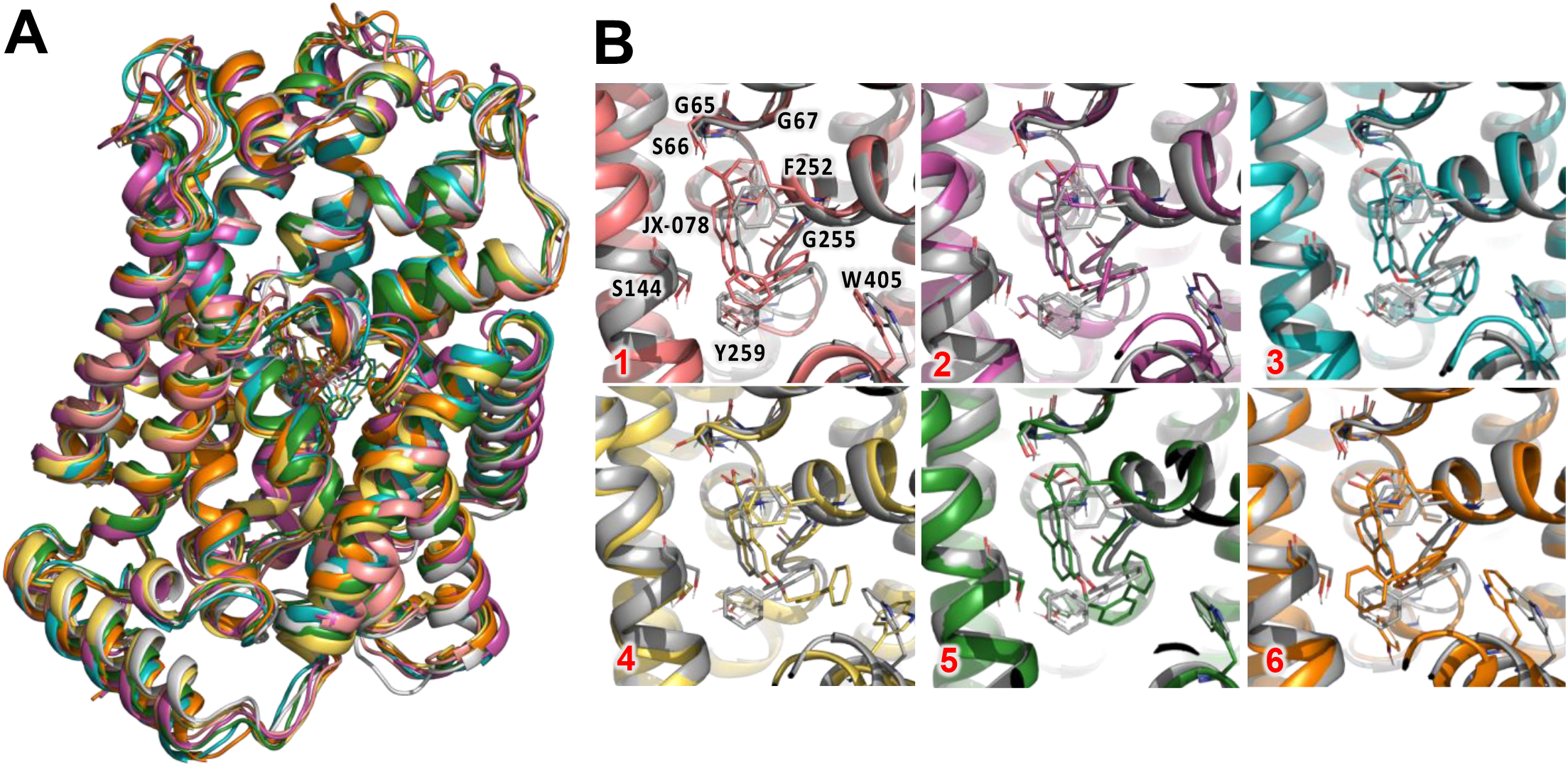
LAT1 structural heterogeneity. Ensemble conformations of LAT1 generated from outward-occluded metainference simulation are shown. **(A)** Superposition of six representative models (clusters) following ensemble metainference simulation of the outward-occluded state. The models display a mostly stable arrangement of the overall structure and ligand binding site. **(B)** Superposition of the outward-occluded, JX-078 bound structure (PDB identifier 7DSL (grey)) and each labeled individual model (colored), focusing on the substrate binding site.

We analyzed the differences and similarities among the models, focusing on the interactions between JX-078 and LAT1 (Fig. 3B). The ligand conformations remained consistent among the representative models, with the amino and carboxy group of JX-078 making sustained hydrogen bonds with the same residues highlighted from our enrichment study, i.e., S66, G67, F252, G255, maintaining hydrogen bonding for over 90% of the simulation (Fig. S2). We also observed that these interactions were further stabilized by Y259 making π-stacking interactions with the biphenyl ring that connects to the amino group (Fig. S2). Additionally, we observed that the hydrophobic tail of JX-078 consistently binds deeper into the binding pocket but explores regions of the binding site, revealing two sub-pockets, C and D (Fig. 5A), Interestingly, we observed that inhibitor showed some preference for sub-pocket C (Fig. 5A), as seen in models 4, 5 and 6, with residue W405 playing a key role in inhibitor binding by making an additional π-stacking interaction with the benzyloxy ring of the inhibitor (Fig. S2). Most of the simulation displayed a binding pose targeting sub-pocket D, in the ensemble simulation, representing over 90% of all the conformations found (Fig. 3B: Models 1 & 6; Fig. S1). We hypothesized that these sub-pockets can be exploited with rational ligand design (Fig. 5E).

We also assessed the stability and movement of the LAT1-JX-078 structure throughout the simulation. We observed that the overall structure and binding site residues remained stable, which is reflected in the low RMSD of all backbone atoms throughout the simulation and particularly in the binding site residues (Fig. 4A). Closer inspection of the individual residues of LAT1 by Root Mean Square Fluctuation (RMSF) analysis, revealed that most of structural variability compared with the cryo-EM structure occurs away from the binding site, with movement less than 1 Å (Fig. 4B). This analysis supports the notion that the size and shape of the LAT1 binding pocket in the outward-occluded conformation remains mostly stable, with most of the flexibility occurring at TM10. The inhibitor displayed some flexibility (Fig. 4C) showing some movement as great as 5 Å. Most of this flexibility stems from the hydrophobic tail of the inhibitor, which is captured by the representative models 1-6 (Fig. 3B). Notedly, the stability of ligand mirrors the increase in contacts between the residues: I63, S65, S66, G67, F252, G255 and Y259. We speculate that these residues are essential to JX-078 binding to LAT1.

**Figure 4:**
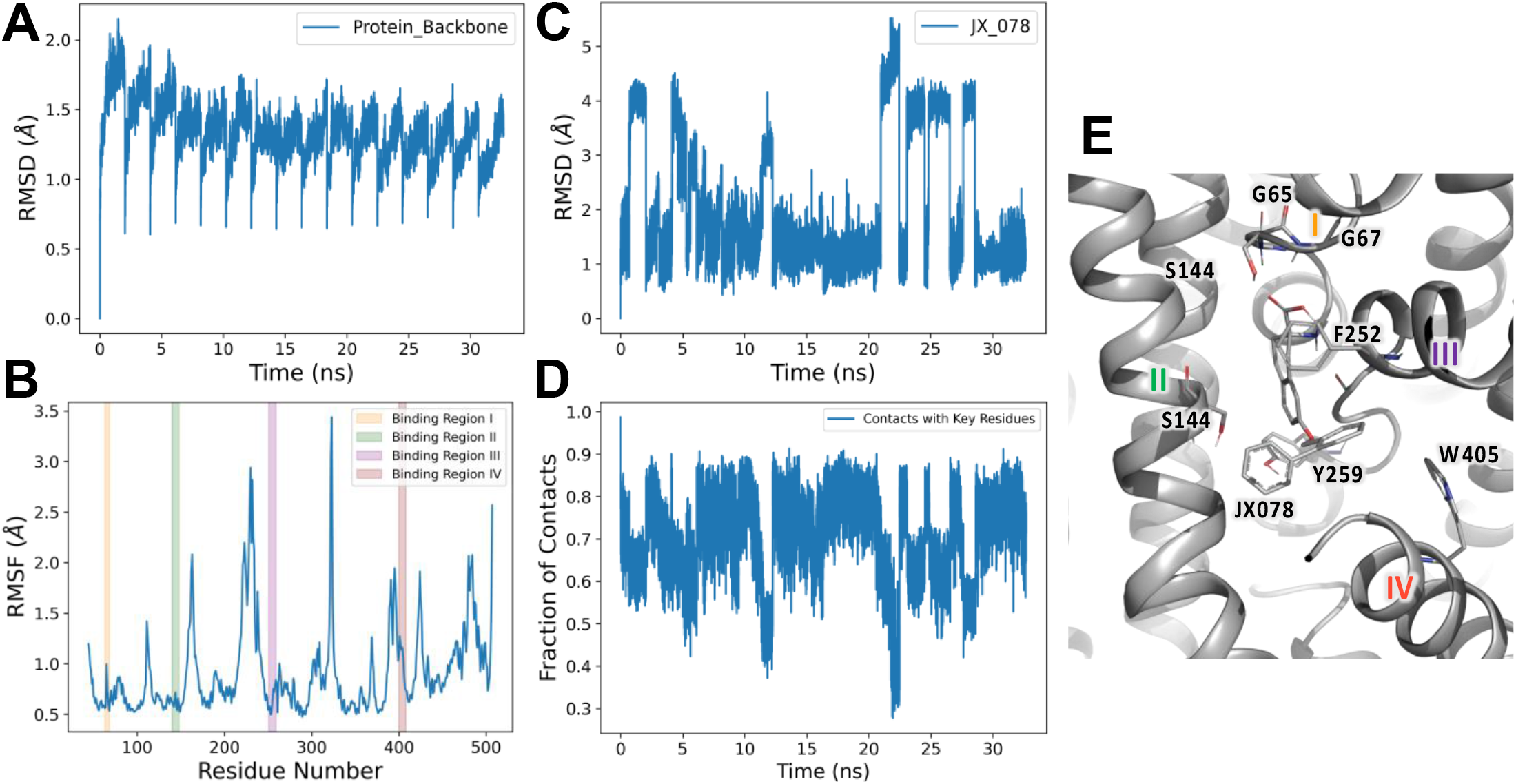
Quantitative analysis of binding site movements. **(A)** Root Mean Square Deviation (RMSD) analysis of metainference simulation focusing on the backbone α-carbon atoms of all residues **(B)** Root Mean Square Fluctuation (RMSF) analysis of individual residues. Binding site residues are broken down into four regions: I, II, II IV in yellow, green, purple, and red, respectively. **(C)** RMSD analysis of JX-078’s atoms. **(D)** Fraction of contacts made between residues (I63, S65, S66, G67, F252, G255, Y259) in the LAT1 substrate binding site and JX-078 throughout the simulation. **(E)** Binding site of 7DSL with each region from **(B)** identified.

### Virtual screening and functional testing of novel compounds for LAT1

Next, we computationally screened a library of purchasable compounds from the ZINC20 lead-like database (3.8 million compounds) and amino-acid like compounds from Enamine (8,137 compounds), against the outward-occluded conformation. The top scoring compounds from each screen were prioritized for further analysis (Methods). Specifically, we prioritized compounds for experimental testing based on the following considerations:

i. Compounds contain both carboxy and amino group that make critical interactions with binding site residues.
ii. Compounds are predicted to make hydrogen bonds with S66, G67, G255 and F255.
iii. Compounds have a hydrophobic tail that extends deeper into the binding site, preferably to sub-pocket C or D (Fig. 5A).

**Figure 5:**
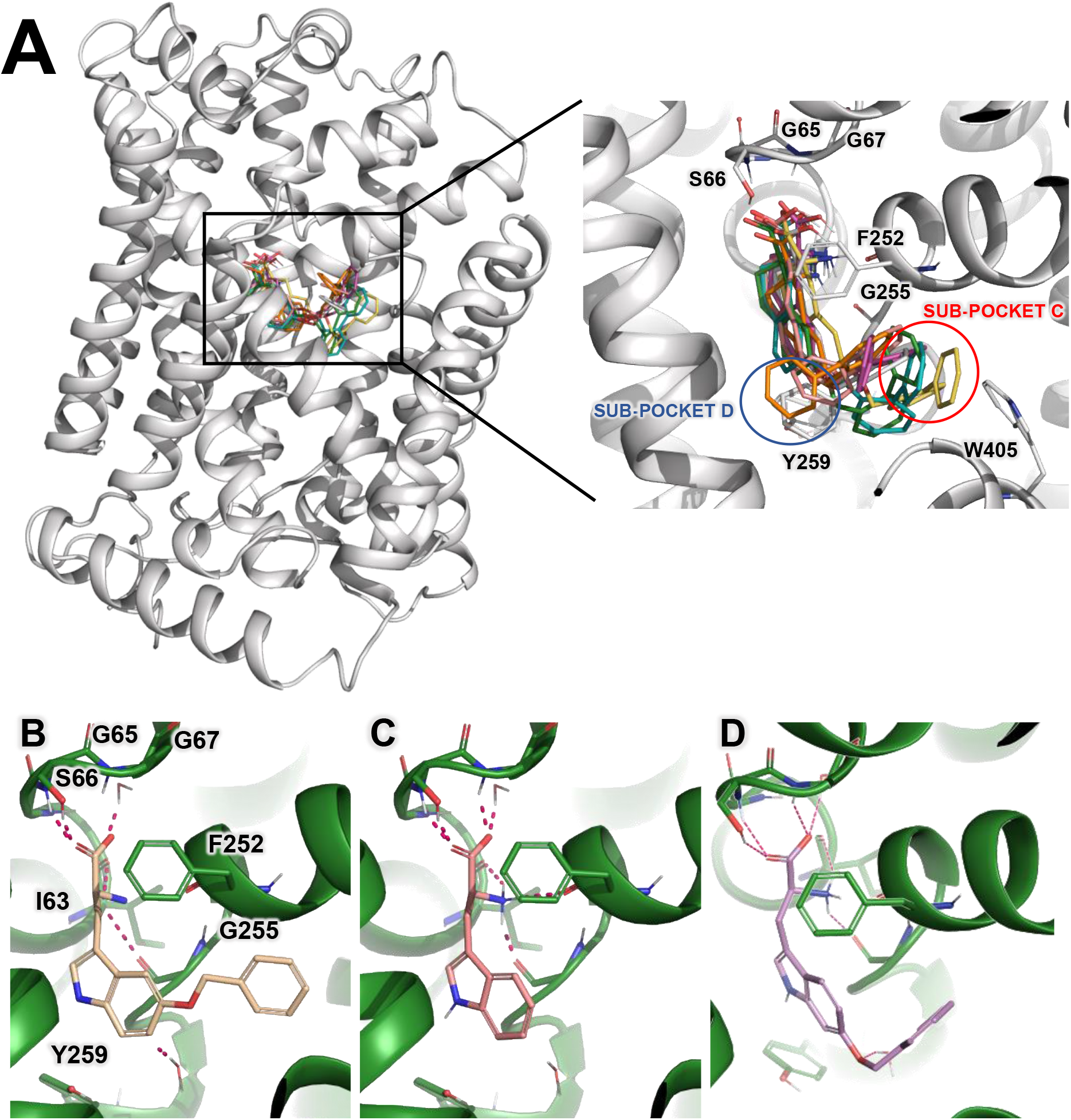
Predicted binding mode of newly discovered LAT1 inhibitors. **(A)** Binding poses of JX-078 to LAT1 derived from metainference simulations using the cryo-EM structure (PDB identifier 7DSL). Predicted key residues from enrichment analysis (Fig. 1). The compound extends to druggable sub-pockets C and D. Docking pose of (**B**) KH-13 and **(C)** tryptophan, which is a known substrate of LAT1. **(D)** Docking pose of a tryptophan analog that targets Pocket C.

**Figure 6:**
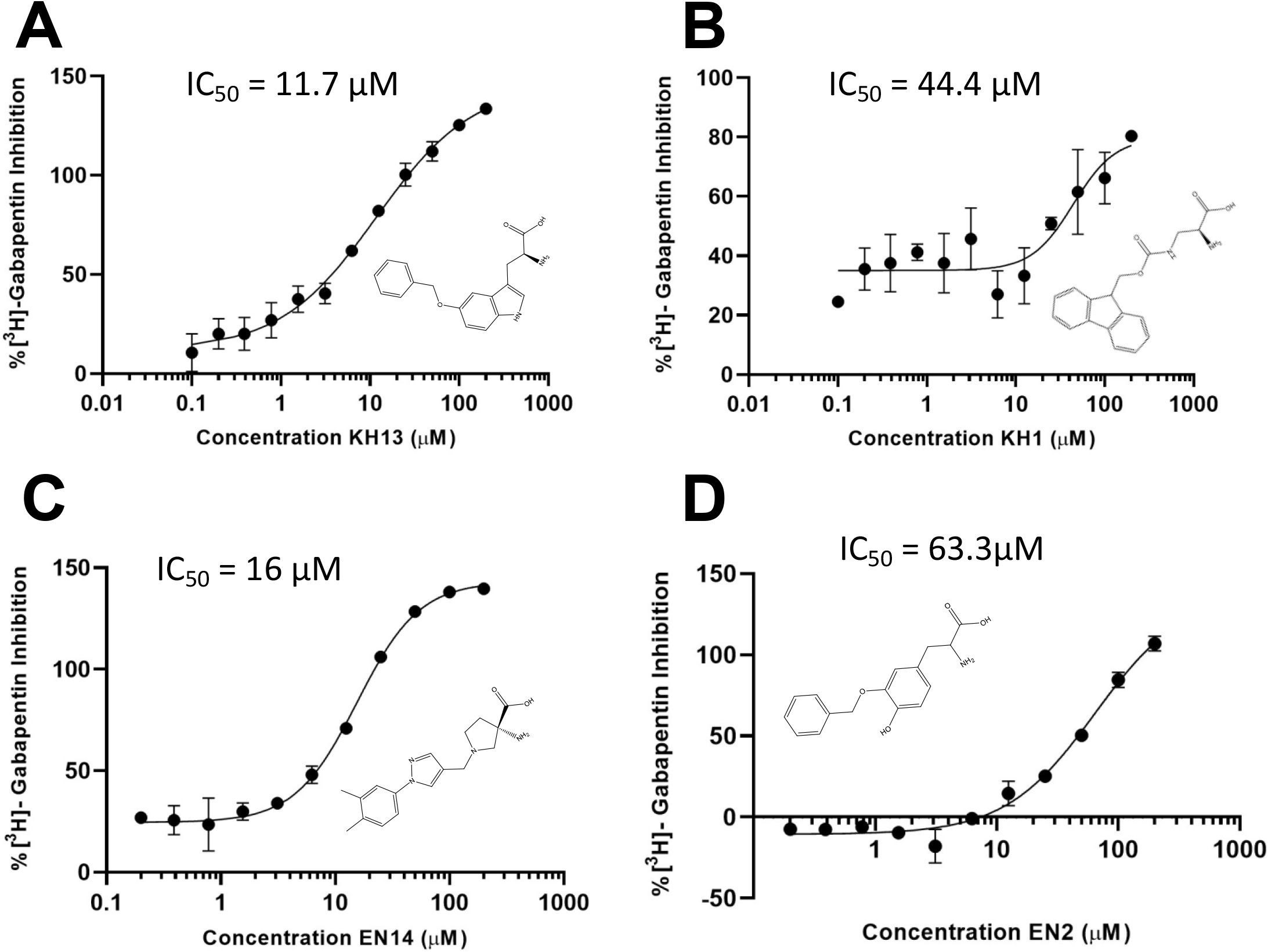
IC_50_ values of newly LAT1 inhibitors. 2-D representation of **(A)** Compound KH13 and **(B)** Compound KH1 **(C)** Compound EN2 **(D)** Compound EN14 and **(E)** L-Phenylalanine, along with each representative dose-dependent inhibition curve. For IC_50_ determination, varying concentrations of each compound were tested ranging from 0.1 μM to 200 μM. GraphPad Prism version 9.3 was used to calculate IC_50_ values, and each concentration point (n=2) was normalized to L-Phenylalanine at 200 μM.

Overall, 24 compounds were initially selected for experimental testing using a *cis*-inhibition assay, which is based on an inhibitor’s capacity to reduce intracellular accumulation of the probe substrate [^3^H]-gabapentin using stable TREx HEK-hLAT1 cell line (see materials and methods). Eleven compounds were identified as LAT1 inhibitors by reducing cellular uptake by > 50% at inhibitor concentration of μM 200 (Table S1). The most potent compound was **KH13** (Fig. 5B), which exhibited an IC_50_ value of 11.8 μM. **KH13** is a benzyloxy-substituted tryptophan, and its predicted binding pose is nearly identical to that of tryptophan (Figs. 5B and 5C). the benzyloxy group of **KH13** interacts with the newly discovered sub-pocket C and W405 and likely contributes to its potency. We designed and tested analogs of **KH13**, three benzyloxy regioisomers and two benzyl amides (**AT182-186**), that were targeted to the same sub-pockets (*e.g*., **AT185**, Fig. 5D). These analogs exhibited comparable potency to that of **KH13** (Table S1). These results increased our confidence in the predicted binding mode of our initial inhibitors. Other new inhibitors included the phenylalanine analog **EN14** (IC_50_ = 16 μM) and tyrosine analog **EN2** (IC_50_ = 63.3 μM). **EN14** contains a unique diazo linkage, and like **KH13**, it places a hydrophobic phenyl ring into sub-pocket C and is also a potent inhibitor. Interestingly, the benzyloxy group in **EN2** docks into sub-pocket D, and the decrease in potency relative to **KH13** suggests this is a less optimal position for the benzyloxy group. Finally, **KH1** (IC_50_ = 44.4 μM) contains a bulky fluorenylmethoxycarbonyl (Fmoc) sidechain that interacts with both sub-pockets C and D. Taken together, these newly discovered LAT1 inhibitors represent diverse scaffolds and/or novel binding modes.

## DISCUSSION

LAT1 is physiologically important protein and a key drug target for a range of diseases and disorders. Despite significant improvement in our understanding of LAT1 structure and function over the past decade, the description of LAT1-ligand interactions at the molecular level is still inadequate. Structural description of LAT1 conformations and mode of binding with ligands is needed for understanding LAT1 transport, as well as for the development of novel small molecule modulators of its function. The goal of this study was to characterize LAT1-inhibitor interactions by combining computational and experimental approaches. The following three key results emerge from this study.

### Characterizing LAT1’s binding site and ligand binding

First, we identified key binding site residues and propose the binding modes of known and newly discovered inhibitors. Ligand enrichment (Fig. 1) in combination with metainference analysis (Figs. 3,4), highlighted S66, G67, F252 and G255 as critical residues for inhibitor binding in agreement with previous studies^42–46,79^. We also propose the importance of an additional two residues: Y259 and W405. The backbone atoms of S66 and G67 form hydrogen bonds with the JX-078 amino group, and those of F252 and G255 form hydrogen bonds with the inhibitor’s carboxy group (Fig. 1,3). Furthermore, analysis of cryo-EM structures of LAT1 in two different conformations (i.e., outward-occluded and inward-open) in complex with different inhibitors (i.e., BCH and JX-078, respectively) revealed that the interactions between the amino and carboxy moieties of the ligands and LAT1 remain mostly consistent, with only a subtle rearrangement of the backbone residues of the ligand binding site (Fig. 3A). Of note, the binding site in the outward-occluded structure is significantly larger (Fig. 2) and possesses potential sub-pockets that are not seen in the inward-open conformation (Fig. 3,5), in part due to the orientation of the Y259 sidechain (Fig. 2). Our metainference analysis revealed two potential druggable sub-pockets that the hydrophobic tail of JX-078 fits into (Figs. 3,5), with W405 playing a key role in sub-pocket C binding. Overall, our results suggest that the outward-occluded structure is a more relevant conformation for rational drug design than the inward-open conformation.

### Identifying unique LAT1 inhibitors

Second, we identified previously unknown pharmacophores of LAT1 inhibitors. Currently, there is a limited number of small molecule tool compounds targeting LAT1, representing few chemical scaffolds^11,79^. Structure-based virtual screening allows the exploration of novel scaffolds by predicting binding of large compound libraries with minimal bias. Although the newly experimentally validated inhibitors from the virtual screens, including those in this study, are not optimized, and are thus, not sufficiently potent to be lead compounds, some of them possess chemical moieties that can serve as starting points for the exploration of unique chemical spaces. For example, **KH14** contains a distinctive pyrrolidine-pyrazole moiety that offers opportunities to substitute with different groups off the pyrazole ring nitrogen; and **KH1** shows that a carbamate moiety can be used to project a 9H-fluorene ring into both sub-pockets C and D. **EN14** contains an uncommon aryl azo group and extends deep into sub-pocket C and provides an opportunity for additional modification by replacing and/or functionalizing the phenyl ring.

While other newly identified compounds are derived from known scaffolds, these compounds exhibited unique modes of binding. For example, substituted tryptophan **KH13** and substituted tyrosine **EN2** both extend deep into the previously uncharacterized sub-pockets C and D, respectively. While extensive SAR work has been done on investigating phenylalanine^11,43,63^ and tyrosine^43^ derivatives, until now, tryptophan has been largely overlooked as a starting point for inhibitor or prodrug design. Thus, this work provides a framework for developing future tryptophan analogs, by building upon the results of **KH13** and **AT182-186**, all of which are potent and novel LAT1 inhibitors.

### Demonstrating the utility of metainference MD simulations

Third, we demonstrated an application of emerging integrative modeling approach to characterize the transporter binding site and rationally design small molecule ligands. Cryo-EM is emerging as the primary method for structure determination of membrane proteins including SLC transporters^71,72^. However, cryo-EM structures are often not determined in a resolution sufficient for structure-based compound design. Further, this challenge is amplified by the inherent flexibility of membrane transporters, including LAT1. Metainference can account for the heterogeneity in cryo-EM maps as a result from the intrinsic dynamics of these molecules, as well as the errors in the measurement^58–59^. We have previously shown that metainference can identify pharmacologically relevant conformations of an unrelated amino acid transporter and guide the optimizations of inhibitors^60^. Here, we apply metainference to identify putative sub-pockets in the binding site, as well as to estimate the biophysical features of the binding site such as flexibility and accessible volume for ligand binding. Remarkably, metainference-generated structures exhibited more favorable stereochemical quality over the experimentally determined structure (Fig. S1), as well as explored ligand and transporter conformations that guided the identification of unique LAT1 inhibitors. As more structures of LAT1 and other SLCs are solved with cryo-EM, metainference and other computational approaches such as the machine learning based approach AlphaFold2^82,83^, can be applied to identify novel ligands for effective therapy and study of human diseases.

## Author Contributions

K.H., D.B.S, A.S., A.A.T. and M.B. conceptualized and designed the study. K.H. performed molecular docking and virtual screening, supervised by A.S. Metainference MD simulations were performed by K.H. and the results and analysis were reviewed and supervised by M.B. Compound design and synthesis were performed by A.A.T, J.B. and C.C. Experimental functional testing was performed by D.B.S. The manuscript was drafted by K.H. and A.S and A.A.T, M.B., D.B.S, J.B., C.C. contributed to the writing of the final manuscript.

## Acknowledgments

Funding for this work was provided by the following grants: R01 GM108911 (A.S.), T32 GM062754 (K.H.), and R15 NS099981 (A.A.T.)

This work was supported in part through the computational resources and staff expertise provided by Scientific Computing at the Icahn School of Medicine at Mount Sinai.

M.B. acknowledges the support of the French Agence Nationale de la Recherche (ANR), under grant ANR-20-CE45-0002 (project EMMI).

