## Supporting Material for "Describing Inhibitor Specificity for the Amino Acid Transporter LAT1 from Metainference Simulations"

**FIGURE S1:**

A

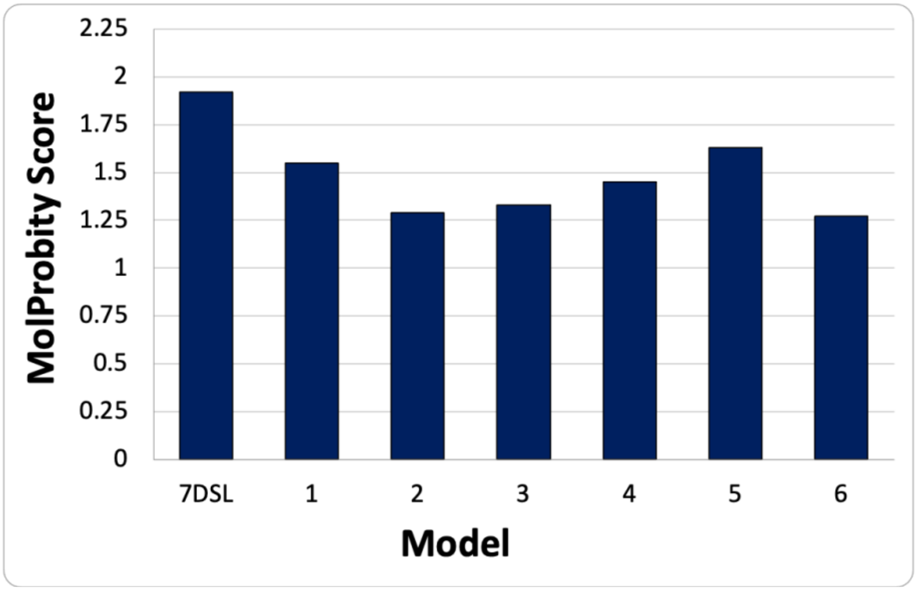


B

| Cluster | Population |
|---|---|
| 1 | 29918 |
| 2 | 1303 |
| 3 | 1130 |
| 4 | 119 |
| 5 | 86 |
| 6 | 80 |

**Fig. S1: MolProbity analysis of Outward-Occluded metainference models. (A)** MolProbity scores of each representative model of each cluster derived from the metainference simulation, compared with outward occluded structure. (PDB Identifier: 7DSL-chain B). **(B)** Table with the number of simulation frames for each cluster the representative models were derived from.

**FIGURE S2:**

A

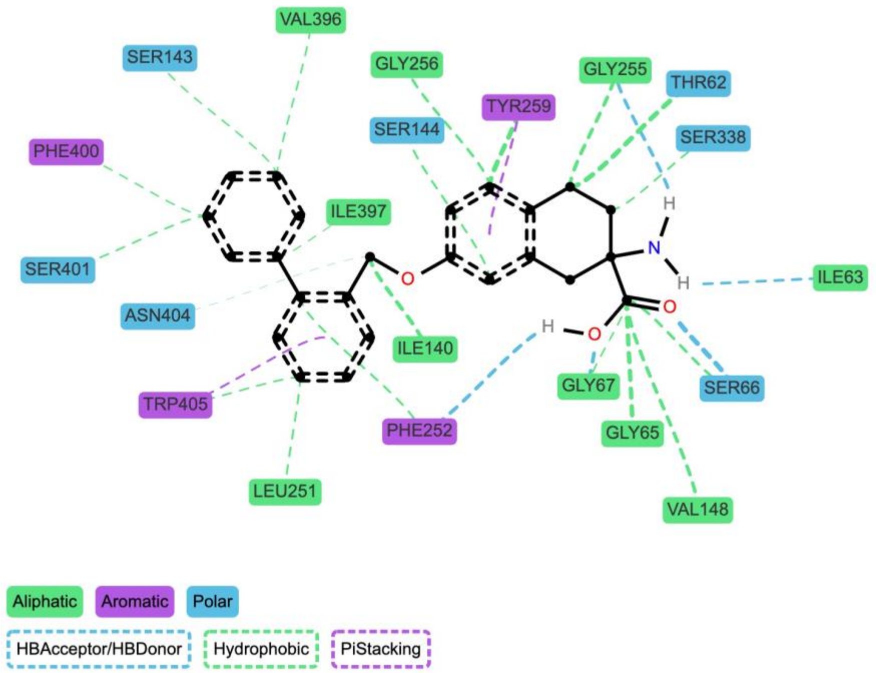


B

| Residue | Interaction Fraction |
|---|---|
| PHE252 | 1.00 |
| GLY255 | 1.00 |
| TYR259 | 0.99 |
| SER66 | 0.94 |
| THR62 | 0.94 |
| GLY67 | 0.93 |
| ILE140 | 0.92 |
| SER144 | 0.92 |
| TRP405 | 0.89 |
| GLY256 | 0.85 |
| GLY65 | 0.84 |
| VAL148 | 0.83 |
| LEU251 | 0.82 |
| VAL396 | 0.72 |
| ILE63 | 0.72 |
| ILE397 | 0.67 |
| SER401 | 0.63 |
| SER338 | 0.48 |
| SER143 | 0.44 |
| ASN404 | 0.39 |
| PHE400 | 0.32 |

**Fig. S2: JX-078 interaction profile. (A)** Interaction map highlighting specific interactions (Hydrogen Bonds, Hydrophobic and π-interactions) between JX-078 and residues in the LAT1 binding site, found during metainference simulation. **(B)** Table displaying the percentage of the sum of all interactions found between individual residues in LAT1 binding site and JX-078 during the entire simulation. (cut-off: 0.3)
